# Wild tobacco genomes reveal the evolution of nicotine biosynthesis

**DOI:** 10.1101/107565

**Authors:** Shuqing Xu, Thomas Brockmöller, Aura Navarro-Quezada, Heiner Kuhl, Klaus Gase, Zhihao Ling, Wenwu Zhou, Christoph Kreitzer, Mario Stanke, Haibao Tang, Eric Lyons, Priyanka Pandey, Shree P. Pandey, Bernd Timmermann, Emmanuel Gaquerel, Ian T. Baldwin

**Author notes:** Correspondence to Dr. Shuqing Xu or Dr. Emmanuel Gaquerel or Prof. Ian T. Baldwin. these authors contributed equally (Shuqing Xu, Thomas Brockmöller).

## Abstract

Nicotine, the signature alkaloid of *Nicotiana* species responsible for the addictive properties of human tobacco smoking, functions as a defensive neurotoxin against attacking herbivores. However, the evolution of the genetic features that contributed to the assembly of the nicotine biosynthetic pathway remains unknown. We sequenced and assembled genomes of two wild tobaccos, *Nicotiana attenuata* (2.5 Gb) and *N. obtusifolia* (1.5 Gb), two ecological models for investigating adaptive traits in nature. We show that after the Solanaceae whole genome triplication event, a repertoire of rapidly expanding transposable elements (TEs) bloated these *Nicotiana* genomes, promoted expression divergences among duplicated genes and contributed to the evolution of herbivory-induced signaling and defenses, including nicotine biosynthesis. The biosynthetic machinery that allows for nicotine synthesis in the roots evolved from the stepwise duplications of two ancient primary metabolic pathways: the polyamine and nicotinic acid dinucleotide (NAD) pathways. While the duplication of the former is shared among several Solanaceous genera which produce polyamine-derived tropane alkaloids, the innovation and efficient production of nicotine in the genus *Nicotiana* required lineage-specific duplications within the NAD pathway and the evolution of root-specific expression of the duplicated Solanaceae-specific ethylene response factor (ERF) that activates the expression of all nicotine biosynthetic genes. Furthermore, TE insertions that incorporated transcription factor binding motifs also likely contributed to the coordinated metabolic flux of the nicotine biosynthetic pathway. Together, these results provide evidence that TEs and gene duplications facilitated the emergence of a key metabolic innovation relevant to plant fitness.

## Significance Statement

Plants produce structurally diverse specialized metabolites, many of which have been exploited in medicine or as pest control agents, while some have been incorporated in our daily lives, such as nicotine. In nature, these metabolites serve complex functions for plants’ ecological adaptation to biotic and abiotic stresses. By analyzing two high-quality wild tobacco genomes, we provide an in-depth genomic study that directly associates genome evolution with the assembly and evolution of the nicotine biosynthetic machinery. These results demonstrate the importance of the interplay of gene duplications and transposable element insertions in the evolution of the multigenic biosynthetic pathways required of specialized metabolism and illuminates on how complex adaptive traits could evolve.

## Introduction

The pyridine alkaloid nicotine, whose addictive properties are well-known to humans, is the signature compound of the genus *Nicotiana* (Solanaceae). In nature, nicotine is arguably one of the most broadly effective plant defense metabolites, in that it poisons acetylcholine receptors and is thereby toxic to all heterotrophs with neuromuscular junctions. Field studies using genetically-modified *N. attenuata* (coyote tobacco) plants, an annual wild diploid native to Western North America, have revealed that this toxin fulfils multifaceted ecological functions that contribute to plant fitness (1, 2). The strong transcriptional up-regulation of the nicotine biosynthetic machinery in roots in response to herbivore attack of the shoot combined with the active translocation and storage of this toxin provides *N. attenuata* plants with an inducible protection mechanism against a broad spectrum of herbivores (2). In addition, the transport and non-homogenous distribution of nicotine in the nectar of flowers within an inflorescence modifies the trap-lining behavior of humming bird pollinators to maximize outcrossing rates (3). These two facets of the ecological utility of nicotine result from the prolific production of this toxin which can accumulate up to 1% of the leaf dry mass in wild tobacco species (4). This prodigious biosynthetic ability is based on an efficient biosynthetic machinery composed of multiple genes co-expressed in roots (5, 6). In contrast to the deep knowledge on nicotine’s biosynthesis and ecological functions, the evolution of genomic features that facilitated the assembly of a pathway so critical for the survival of *Nicotiana* species has remained largely unknown.

Gene duplication and TE insertions continuously shape the evolutionary landscape of genomes and can affect the function of genes with adaptive consequences (7, 8). While whole-genome and local gene duplications provide the raw material for the evolution of novel traits, TE mobility can broadly remodel gene expression by redistributing transcription factor binding sites, shaping epigenetic marks and/or providing target sequences for small regulatory RNAs (7–11). Hence, the combination of gene duplications and TE activity is thought to facilitate the evolution of novel adaptive traits (8). However, the details of this process, in particular its role in the evolution of metabolic complexity through the assembly of novel multi-gene pathways, remains unclear.

## Results and Discussion

### Genome sequencing, assembly and annotation

We sequenced and assembled the genome of *N. attenuata*, using 30x Illumina short reads, 4.5x 454 reads, and 10x PacBio single-molecule long reads. We assembled 2.37 Gb of sequences representing 92% of the expected genome size. We further generated a 50x optical map and a high-density linkage map for super-scaffolding (Fig. S1 and S2), which anchored 825.8 Mb to 12 linkage groups and resulted in a final assembly with a N50 contig equal to 90.4 kb and a scaffold size of 524.5 kb (Fig. S3). Likewise, using ~50x Illumina short reads, we assembled the *N. obtusifolia* genome with a 59.5 kb and 134.1 kb N50 contig and scaffold N50 size, respectively. The combined annotation pipeline integrating both hint-guided AUGUSTUS and MAKER2 gene prediction pipeline predicted 33,449 gene models in the *N.attenuata* genome. More than 71% of these genes models are fully supported by RNA-seq reads and 12,617 and 18,176 of these genes are orthologous to *Arabidopsis* and tomato genes, respectively.

To investigate the evolutionary history of the different *Nicotiana* genomes, we inferred 23,340 homologous groups using protein sequences from 11 published genomes (Table S1). A phylogenomic analysis of the identified homologous groups demonstrated that *Nicotiana* species share a whole-genome triplication (WGT) event with *Solanum* species, such as tomato, potato and *Petunia* (12), but not with *Mimulus* (Figure 1 and Fig. S4-S7). At least 3,499 duplicated gene pairs originating from this WGT were retained in both *Nicotiana* and *Solanum*. Among all retained duplicated gene pairs detected in *N.attenuata* that did not further duplicate in this species, more than 53.7% showed expression divergence (fold change greater than 2) in at least one tissue, indicating that these WGT-derived duplicated genes may have evolved divergent functions through neofunctionalization or sub-functionalization.

**Figure 1.**
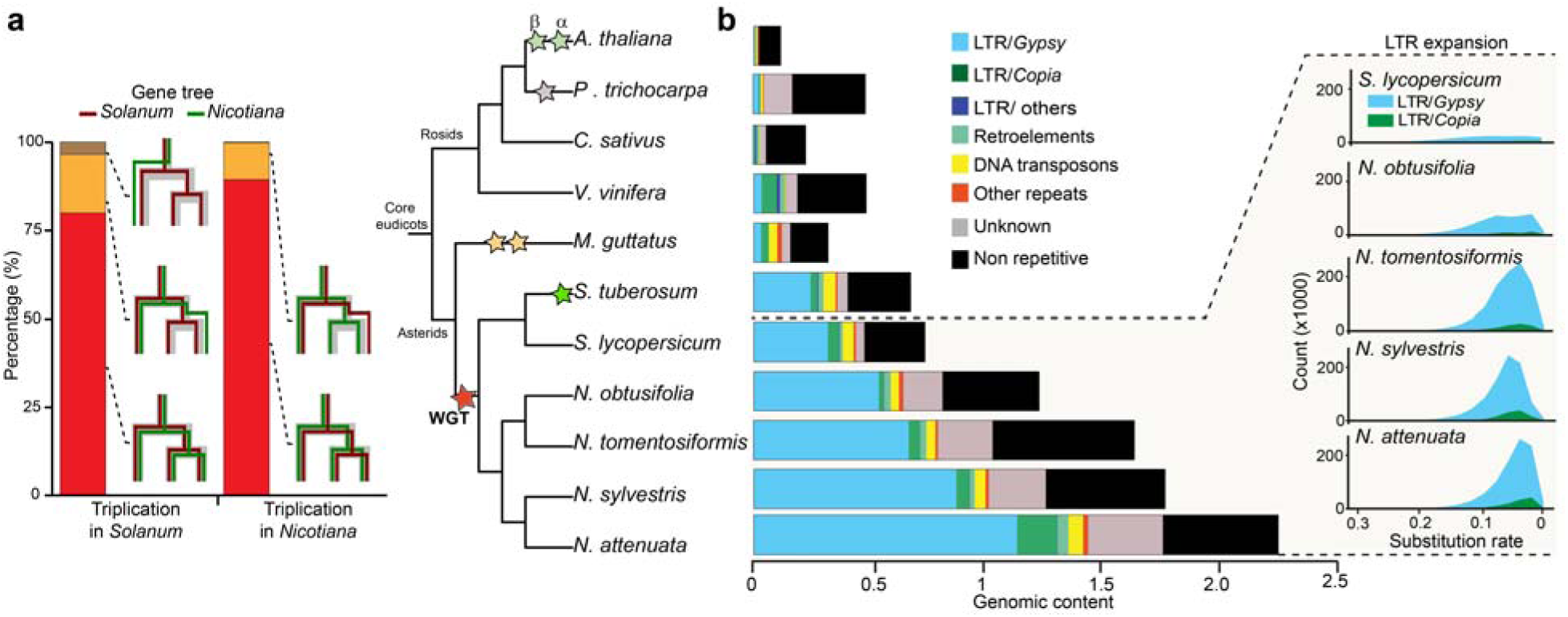
Whole genome triplication (WGT) in *Nicotiana* genomes is shared with other Solanaceae species but the *Gypsy* retrotransposons expansions are *Nicotiana*-specific. (a) *Nicotiana* genomes share the WGT with other Solanaceae species. Left panel depicts the shared WGT event between *Nicotiana* and *Solanum* as revealed by the structure of the phylogenetic tree of triplicated gene families in *Nicotiana* and *Solanum*. Red and yellow bars represent the percentage of triplication and duplication events shared between *Nicotiana* and *Solanum*, respectively. Right panel shows the phylogenetic tree of 11 plant species and different colored stars indicate previously characterized whole genome multiplication events. (b) Expansion of *Gypsy* transposable elements contributes substantially to genome size evolution in *Nicotiana*. Left panel shows the genomic content (in Gb) of repetitive versus non-repetitive sequences in the 11 plant genomes. Black and grey bars indicate non-repetitive sequences, whereas other colors indicate repetitive sequences. The right insert visualizes the expansion history of LTR retrotransposons in four *Nicotiana* genomes in comparison to tomato. X-axis (number of substitutions per site) refers to the divergence of a LTR from its closest paralog in the genome, with smaller numbers indicating more recent duplication events.

### Expansion of transposable elements in *Nicotiana*

Polyploidization is often associated with a burst of TE activity as a hypothesized consequence of “genomic shock” (13, 14). TEs, especially long terminal repeats (LTRs) are highly abundant in *Nicotiana* and account for 81.0% and 64.8% of the *N. attenuata* and *N. obtusifolia* genomes, representing significantly higher proportions than other sequenced Solanaceae genomes, such as tomato and potato (Figure 1). An analysis of the history of TE insertions revealed that all *Nicotiana* species experienced a recent wave of *Gypsy* retrotransposon expansion. However this expansion of *Gypsy* copies was less pronounced in *N. obtusifolia* compared to other *Nicotiana* species analyzed, which accounts for the smaller genome size of *N. obtusifolia*. A recent study showed that *Capsicum* species also experienced a large expansion of their *Gypsy* repertoire (15), albeit earlier than in *Nicotiana*, indicating that after WGT, the different Solanaceae lineages independently experienced the processes of *Gypsy* proliferation.

In addition to LTRs, miniature inverted–repeat transposable elements (MITEs), which are derived from truncated autonomous DNA transposons, may also play evolutionary roles. Although the size of MITEs is generally small, typically less than 600 bp, MITEs are often located adjacent to genes and are often transcriptionally active. As such, they have been hypothesized to contribute to the evolution of gene regulation (16, 17). In total, we identified 13 MITE families in the genome of *N. attenuata*, several of them having rapidly and specifically expanded in *Nicotiana* species (Figure 2a and b). Among these expanded MITE families, a Solanaceae-specific subgroup of the Tc1/Mariner defined by DTT-NIC1 is the most abundant. By analyzing insertion positions of this subgroup, we found that DTT-NIC1 copies, similar to other DNA transposons, are significantly enriched within a 1 kb region upstream of the genes in *N. attenuata* (Figure 2c). Analyses on the herbivory-induced conserved transcriptomic responses in *Nicotiana* further showed that insertions of DTT-NIC1 are significantly enriched within the 1 kb upstream region of herbivory-induced early defense signaling genes in *N. attenuata*, and may have contributed to the recruitment of genes into the induced defense signaling network by introducing WRKY transcription factor binding sites (18).

**Figure 2.**
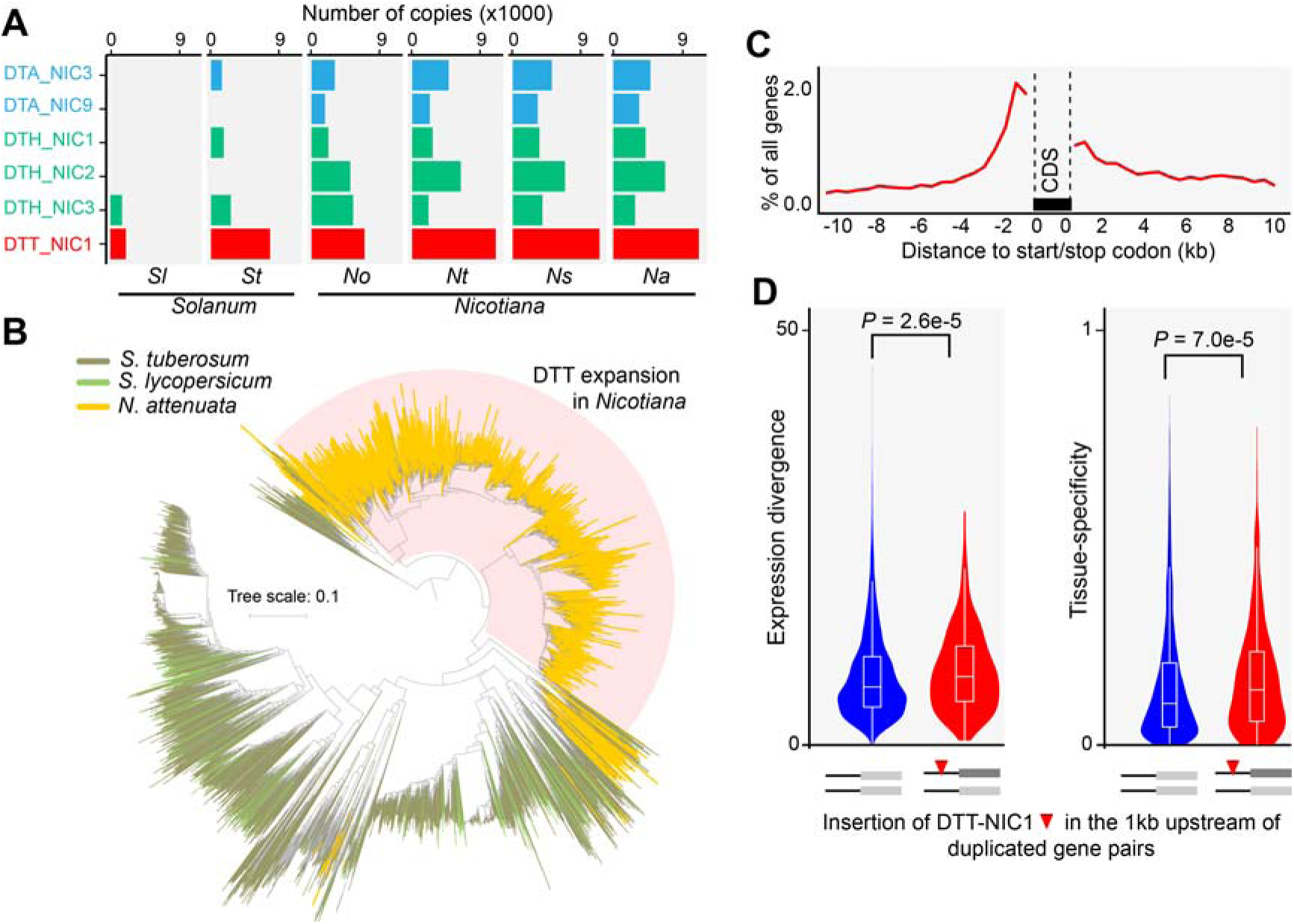
Expansion of transposable elements of the family DTT-NIC1 increased genome-wide gene expression divergence among duplicated gene pairs in *Nicotiana*. (a) Copy number of the six most abundant *Nicotiana* MITE families of transposable elements in *Nicotiana* and *Solanum*. Each bar depicts the total number of copies in each species for the six main MITE transposable elements (TEs). MITE families are visualized by different colours: light blue, DTA (*h*AT); green, DTH (PIF/Harbinger); red, DTT (Tc1/Mariner). DTT-NIC1 from the Tc1/Mariner family is the most abundant all MITE TEs. *Nicotiana* species: *Na*, *N. attenuata*; *No*, *N. obtusifolia*; *Ns*, *N. sylvestris*; *Nt*, *N. tomentosiformis*. *Solanum* species: *Sl*, *Solanum lycopersicum*; *St*, *Solanum tuberosum*. (b) Expansion of the DTT-NIC1 family in *Nicotiana* species. Neighbor joining (NJ) tree of the DTT-NIC1 family in *N. attenuata*, tomato and potato. The shaded clade highlights the pronounced expansion of DTT-NIC1 in *N. attenuata*. (c) DTT-NIC1 insertions are enriched in the upstream regions of coding sequences. The line indicates the percentage of genes, among all predicted protein coding genes, that contain DTT-NIC1 insertions within a given 500 bp sliding window. (d) Insertions of DTT-NIC1 within the 1 kb upstream region of duplicated genes increased tissue-level gene expression divergence (Wilcoxon rank sum test). Left and right panels are violin plots of the divergences between duplicated gene pairs at expression and tissue specificity levels, respectively. Red bars indicate duplicated pairs, of which one copy has at least one DTT-NIC1 insertion and the other does not. Blue bars indicate duplicated pairs, both of which lack DTT-NIC1 insertions. The width of the probability density in the violin plots along the bars correspond to the number of duplicate gene pairs.

Innovations in metabolic and signaling network architecture are thought to result from the rapid rewiring of tissue-level gene expression patterns following duplications events (19, 20). To examine this inference, we compared the genome-wide expression divergence between duplicated gene pairs and analyzed the effects of DTT-NIC1 insertions into 1 kb upstream regions of each member of the gene pairs. Insertions of the DTT-NIC1 family were associated with significant divergences in expression and tissue specificity between duplicated genes (Figure 2d), consistent with the hypothesis that the expansion of this TE family was a critical determinant of genome-wide re-wirings of gene regulation occurring post-duplication in these *Nicotiana* species.

### Evolution of nicotine biosynthesis

To further understand the role of gene duplication and TE insertions on the evolution of *Nicotiana* adaptive traits, we reconstructed the evolutionary history of the nicotine biosynthesis pathway, a key defensive innovation of the *Nicotiana* genus. Nicotine biosynthesis is restricted to the roots and involves the synthesis of a pyridine ring and a pyrrolidine ring which are coupled most likely via the action of genes coding for an isoflavone reductase-like protein, called A622, and the berberine bridge enzyme-like (BBL) enzymes (21, 22) (Figure 3A). Phylogenomic analyses revealed that genes involved in the biosynthesis of the pyridine and pyrrolidine rings evolved from the duplication of two primary metabolic pathways that are ancient across all plant lineages: the nicotinamide adenine dinucleotide (NAD) cofactor and polyamine metabolism pathways, respectively (Figure 3a).

**Figure 3.**
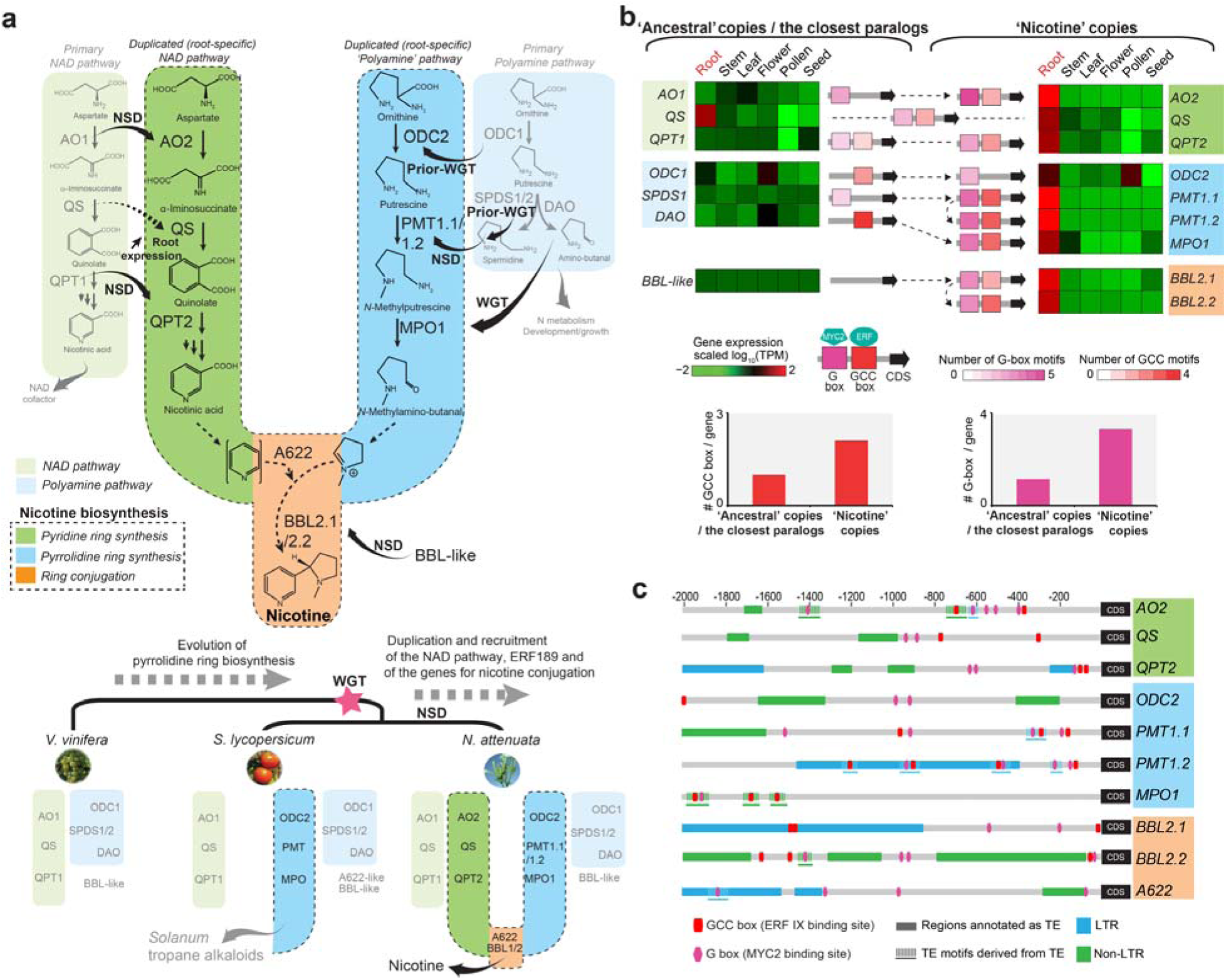
Prolific nicotine production evolved from the duplication of two primary metabolic pathways and its coordinated transcriptional regulation was likely facilitated by transposon-derived transcription factor binding site insertions. (a) *Nicotiana*-specific nicotine biosynthesis originates from step-wise duplications of two primary metabolic pathways. The upper panel depicts the metabolic organization (brightly colored and dashed line outlined branches) and evolution of nicotine biosynthesis via pathway and single gene duplications in *Nicotiana*. Light green and light blue branches on the side indicate the two ancient gene modules with housekeeping functions in plants corresponding to the NAD cofactor and polyamine pathways. Different gene duplication types are indicated by arrows annotated as follows: NSD, *Nicotiana*-specific duplications; WGT, whole genome triplication in Solanaceae; Prior-WGT, gene duplication occurring prior to WGT. *Nicotiana* QS did not duplicate but experienced an increase in root expression compared to its tomato homolog. Lower panel: phylogenomics view of grape, tomato and *N. attenuata* gene sets highlighting the gradual assembly of the nicotine biosynthetic pathway. (b) Acquisition of transcription factor binding motifs and root-specific expression evolution of nicotine biosynthesis genes. Heatmaps depict the scaled expression of nicotine biosynthetic genes and their ancestral copies or closest paralogs across six distinct tissues. Red and green signify high and low expression, respectively. A622 likely neofunctionalized without being duplicated in *solanaceous* species. TPM: transcript per million. Nicotine biosynthetic genes’ root specific expression and dramatic transcriptional up-regulation during insect herbivory is coordinated by the action of MYC2 and ERF transcription factors which target G- and GCC-type boxes in the promoters, respectively. Numbers of GCC and G-box motifs detected within 2 kb upstream region of nicotine biosynthetic genes and their ancestral copies are represented using specific color gradients. GCC motifs derived from TE insertion in the gene upstream region are shown as blue lines. (c) Many GCC and G-box motifs from nicotine biosynthesis genes are likely derived from TE. Each row depicts the motif and TE annotation of the 2 kb upstream region of an individual nicotine biosynthesis gene. The predicted GCC and G-box motifs are shown in red and pink small boxes, respectively. The regions that were annotated as TE from RepeatMasker are shown in rectangle with two different colors. Light blue: LTR; green: non-LTR. The motifs sequences and their 150 bp flanking region showed significant homology (E-value less than 1e-5) to annotated TE sequences in *N. attenuata* are shown in dashed lines.

However, the timing and mode of duplications of these two pathways differ and reflect the expansion and recruitment of gene sets required for the diversification of alkaloid metabolism in the Solanaceae. Duplications that gave rise to the branch extension of the polyamine pathway required for the biosynthesis of the signature alkaloids of Solanaceae and Convolvulaceae (e.g., tropane in many genera and nicotine in *Nicotiana*) are shared among *Nicotiana*, *Solanum*, and *Petunia* with individual gene members recruited from the Solanaceae WGT or earlier duplication events. Genes encoding ornithine decarboxylase (ODC2) and *N*-putrescine methyltransferase (PMT) duplicated prior to the shared Solanaceae WGT from their ancestral copies in polyamine metabolism, *ODC1* and *spermidine synthase* (*SPDS*), respectively (Fig. S8 and S9). While ODC2 likely retained its ancestral enzymatic function, PMT (derived from SPDS) acquired the capacity to methylate putrescine to form *N*-methyl-putrescine through neofunctionalization (23). The *N-*methylputrescine oxidase (*MPO*) from the polyamine metabolism pathway evolved from diamine oxidase (DAO)(24) through whole genome multiplication. Both copies were retained in *Nicotiana*, *Solanum* and *Petunia* (Fig. S10), presumably to sustain the flux of *N*-methyl-Δ^1^-pyrrolinium required for alkaloid biosynthesis. Duplication patterns of *ODC*, *PMT* and *MPO* therefore support the ancient origin of the ornithine-derived *N*-methyl-Δ^1^-pyrrolinium, which is utilized as a common building block for the biosynthesis of most alkaloid groups in the Solanaceae and Convolvulaceae.

In contrast to the relatively ancient origin of pyrrolidine ring biosynthesis, duplications of the NAD pathway genes, encoding aspartate oxidase (AO) and quinolinic acid phosphoribosyl transferase (QPT), responsible for pyridine ring biosynthesis are *Nicotiana*-specific and likely occurred through local duplication events (Fig. S11 and S12). BBLs are thought to be involved in the late oxidation step in nicotine biosynthesis that couples the pyridine and pyrrolidine rings, and therefore constitute a key innovation in the *Nicotiana*-specific synthesis of pyridine alkaloids. *BBLs* exhibiting clear root expression specificity and likely evolved through neofunctionalization after gene duplications (Fig. S13).

Tissue-level RNA-seq transcriptome analyses in *N. attenuata* confirmed that while ancestral copies exhibit diverse expression patterns among different tissues, all of the duplicated gene copies recruited for nicotine biosynthesis are specifically expressed in roots (Figure 3b) and also specifically transcriptionally up-regulated in response to herbivory via the jasmonate signaling pathway (25). Experimental work has shown that the transcription factors of the ethylene response factor (*ERF189*) subfamily IX and *MYC2*, play central roles in the up-regulation of nicotine genes(5). Analyzing the evolutionary history of *MYC2* revealed that this gene duplicated at the base of the Solanaceae via genome-wide multiplication or segmental duplications, and both duplicated copies were retained in the genomes of *Nicotiana* and several Solanaceae species (Fig. S14). Interestingly, *ERF189* is located within an ERF cluster (6, 26) as a result of ancestral tandem duplications shared among *Nicotiana*, *Solanum*, *Capsicum* and *Petunia* (Figure 4). After the split between *Nicotiana* and *Solanum*, *ERF189* was further tandemly duplicated independently in each linage, but only evolved root-specific expressions in the diploid species of *Nicotiana* (Figure 4). Because of its essential role in regulating the expression of all nicotine genes(6), the acquisition of a root-specific expression by *ERF189* in the ancestor of *Nicotiana* species likely played a critical role for the coordinated root expression of nicotine biosynthesis genes.

**Figure 4.**
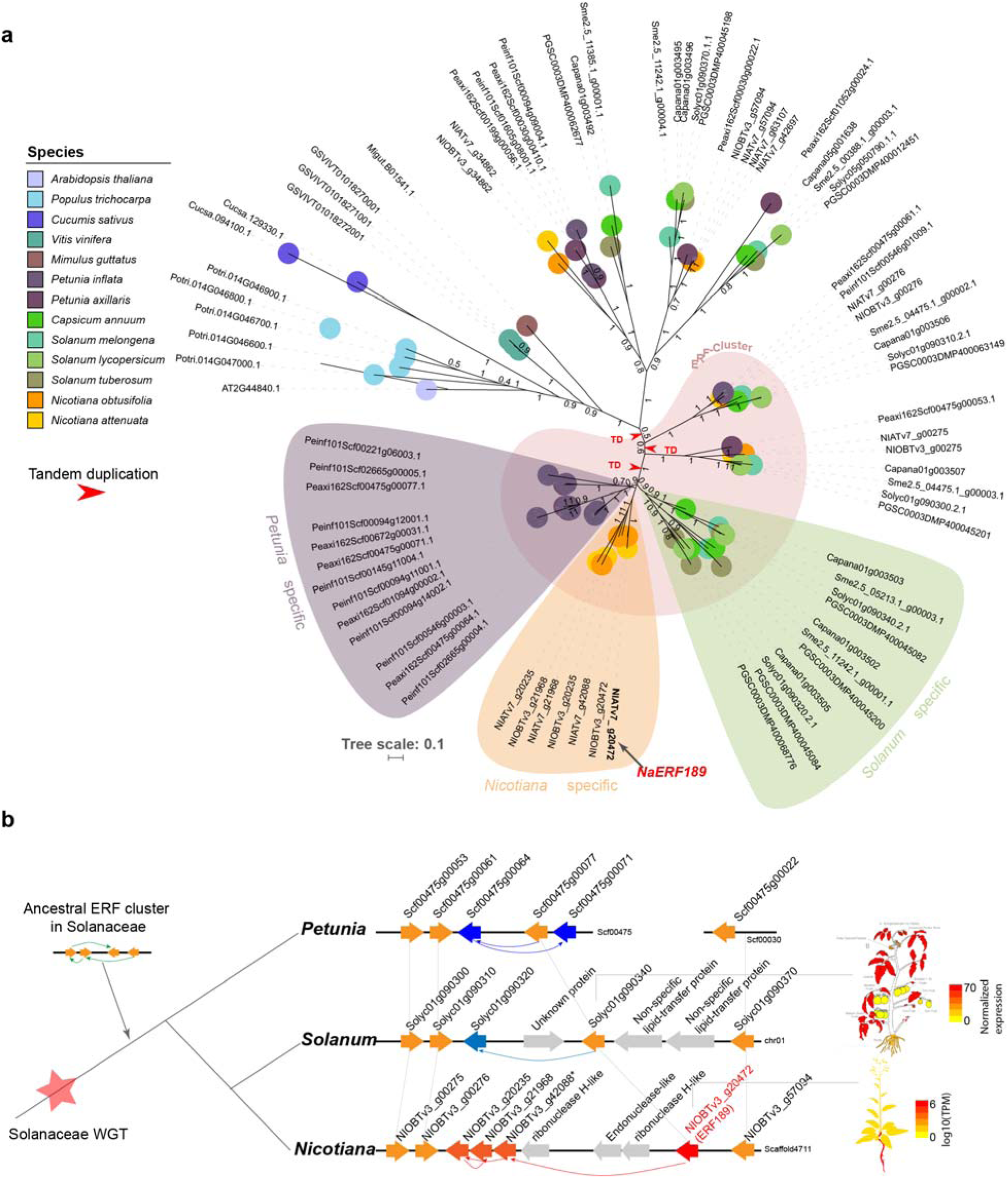
Evolution of *ERF189* in *Nicotiana*. (a) Phylogenetic tree of *ERF189* among different plants. The numbers on the branches denote approximate Bayes branch support values. Red arrows indicate the tandem duplications that were identified based on synteny information. (b) Evolutionary model of the *ERF189* gene cluster based on both phylogenetic relationship among homolog genes and their syntenic information in *P. axillaris* (12), tomato (26) and *N. obtusifolia*. Lineage specific duplications of ERFs are shown in blue (*Petunia*), light blue (*Solanum*) and red (*Nicotiana*). The expression profile of *ERF189* and its orthologous gene in tomato were retrieved from *N. attenuata* data hub (nadh.ice.mpg.de) and tomato eFP browser (http://bar.utoronto.ca/efp_tomato/cgi-bin/efpWeb.cgi), respectively. The asterisk indicates the gene NIOBTv3_g42088 is likely a pseudogene in *N. obtusifolia*.

Regulation of nicotine genes by *MYC2* and *ERF189* relies in part on the presence of two transcription factor binding sites, the GCC and G-box elements in their promotor regions (5, 6, 27). Nicotine biosynthetic genes harbor more than twice the frequency of GCC and G-box elements in their 2 kb upstream region than do their ancestral copies (Figure 3b), consistent with the hypothesis that the accumulation of GCC and G-box elements in promoter regions contributed to the evolution of the coordinated transcriptional regulation required for high-flux nicotine biosynthesis. Investigating the origin of GCC and G-box motifs in upstream regions of 10 nicotine biosynthesis genes showed that at least 34.8% and 29.0% of GCC and G-box motifs, respectively (Figure 3c), are likely derived from TE insertions. While it is unclear whether all of these TE-derived GCC and G-box motifs are involved in regulating the expression of nicotine genes, some likely are. For example, in the case of *PMT1*, a previous experimental study revealed that the 650 bp upstream region, which specifically contained additional TE-derived GCC motifs, had a much larger capacity to drive the expression of a reporter gene in *Nicotiana* roots than did the 111 bp upstream region that lacked these motifs (28). Furthermore, all GCC and G-box motifs within 2kb 5’ region of the *MPO1* that is likely under control of *ERF189* (24) are derived from TEs (Figure 3c).

The mechanisms of genome organizational evolution, such as genome-wide duplications and TE expansions, facilitated the evolution of several aspects of the anti-herbivore defense arsenal including a key metabolic innovation in *Nicotiana* species. These results are consistent with the hypothesis that TEs, which have often been considered as ‘junk’ DNA, can be important orchestrators of the gene expression remodeling that is required for the evolution of adaptive traits. Since native *Nicotiana* species do not survive in nature without the ability to produce large quantities of nicotine to ward off attackers, it is likely that this ‘junk’ has inspired innovation essential for their survival (29).

## Conclusion

We sequenced, *de novo* assembled, and annotated the two genomes of two species of *Nicotiana*, the genus of which are scientifically and economically important. The fully annotated gene models, transposable elements, smRNAs and transcriptomic atlas (*SI Appendix*
Table S1-S5, Dataset S1-S9) of *N.attenuata* enable comparative analysis to illuminate the evolution of specialized metabolites and novel adaptive traits in Solanaceae as demonstrated by our in-depth genomic analysis on the evolution of nicotine biosynthesis.

## Materials and Methods

### Plant material and DNA preparation

Plants were grown as previously described (30). The genomic DNA sequenced by 454 and Illumina HiSeq2000 technologies was isolated from late rosette-stage plants using the CTAB-method(31). The two *N. attenuata* DNA plants used for this extraction were from a 30th generation inbred line, referred to as the UT accession, which originated from a 1996 collection from a native population in Washington County, Utah, USA (30). *N. obtusifolia* DNA was obtained from a single plant of the first inbred generation derived from seeds collected from a native population in 2004 at the Lytle Ranch Preserve, Saint George, Utah, USA. High molecular weight genomic DNA used to generate the optical map of *N. attenuata* was isolated using a nuclei based protocol (32) from approximately one hundred freshly harvested *N. attenuata* plants (harvested 29 days post germination) of the same inbred generation and origin as used for the short-read sequencing.

To reduce the potential effects of secondary metabolites on single molecular sequencing, the plant material used for PacBio sequencing was from a cross of two isogenic *N. attenuata* transgenic lines (mother: ir-*aoc*, line A-07-457-1, which was transformed with pRESC5AOC [GenBank KX011463] and is impaired in JA biosynthesis; father: ir*GGPPS*, line A-07-230-5, which was transformed with pRESC5GGPPS [GenBank KX011462] and is impaired in the synthesis of the abundant 17-hydroxygeranyllinalool diterpene glycosides), both generated from the 22nd inbred generation of the same origin as the *N. attenuata* plants described above (30). Genomic DNA was isolated from approximately one hundred young plants (harvested 29 days post germination) by Amplicon Express (http://ampliconexpress.com) according to a proprietary protocol.

### Genome sequencing and assembly

For *N. attenuata*, the Illumina HiSeq2000 system was used to generate a high coverage whole genome shotgun sequencing (WGS) of the genome based on short reads (2 × 100 bp or 2 × 120 bp). Different paired-end libraries were constructed using the Illumina TruSeq DNA sample preparation kit v2. The fragment size distribution maxima were observed at 180, 250, 600 and 950 bp. Additionally, two mate-pair libraries were constructed using Illumina mate-pair library preparation kit v2, which had their maxima at 5,500 and 20,000 bp of the fragment size frequency distribution, respectively. A lower genome-wide coverage of long reads (median read length 780 bp) was generated by the Roche/454 GS FLX (+) pyro-sequencing technology using Roche rapid library prep kit v2. For *N. obtusifolia*, two paired-end libraries and a single mate-pair library were constructed using the same material. The two paired-end libraries had fragments size distribution maxima at 480 and 1050 bp, respectively. The mate-pair library had a maximum at 3500 bp. The 20 kb mate-pair libraries were constructed at Eurofins/MWG using the Cre-recombinase circularization approach from Roche (Roche Diagnostics GmbH, Mannheim, Germany). These were both sequenced with 454 technology and Illumina HiSeq2000 (by removing Roche adaptor sequences and replacing them by Illumina adaptor sequences). The PacBio reads were sequenced at the Cold Spring Harbor Laboratory.

The overall assembly workflow for *N. attenuata* is shown as Fig. S3. All paired-end reads from the sequenced libraries were assembled using the Celera Assembler (CA7) with a minimum read length cut-off at 64bp. Preliminary tests showed that single end or short reads did not improve the assemblies, but increased calculation time significantly. In total, 86.4 Gb short read data were assembled. The expected genome size of *N. attenuata* is of 2.54 Gb based on coverage of the “larger than N50 length unitigs”, similar to the estimation (1C=2.5pg) from the flow cytometry analysis. We used the SSPACE v2 scaffolder to further improve scaffolding using the mate-pair data and filled gaps using GapCloser v1.12 (33). After manual inspections, we found that certain neighboring contigs in the scaffolds still contained overlaps, which might be due to the assembly process from CA7 that places copies of repeat sequence at the end of contigs or due to issues in Gapcloser v1.12 that leave open some closable gaps. To close these gaps between overlapped contigs, we compared neighboring contigs using BLASTN (min. identity 95%/min. length 43) and then joined overlapping contigs with custom scripts.

The assembly scaffolds from short reads were used for gap filling and further scaffolding using PBJelly (v15.8.24)(34) with ~10x PacBio reads (N50=14.9 kb, max read length =48.9 kb), which resulted in a 2.17 Gb genome assembly. While PBjelly only increased the N50 scaffold size from 176 kb to 202 kb, it significantly increased the N50 contig size from 67 kb to 90 kb. Because PacBio reads contain about 12-15% errors, we performed an additional correction step using short reads with PILON(35). In total, 98.2% of the draft assembly was confirmed by short reads and ~1.3 Mb sequences were corrected by PILON. The PILON corrected assembly was further mapped to the 10x PacBio reads using BLASR and used SSPACE-longreads for the second round scaffolding, which increased the N50 scaffold size to 292 kb.

To assemble the *N. obtusifolia* genome, we employed a hybrid strategy in which we first assembled all short reads by a ‘de Bruijn graph’ assembler using idba-ud v1.1.1 (36), and then assembled the locally re-assembled contigs and a subset of the short read data by an ‘OLC’ long read assembler using CA7. Scaffolding and gap filling were carried out using SSPACE v2 using mate-pair data in a similar manner to the *N. attenuata* assembly.

Analyses of 248 conserved core eukaryotic genes using the CEGMA(37) pipeline indicated that both *N. attenuata* and *N. obtusifolia* assemblies were only slightly less complete in full-length gene contents than that of the tomato genome (86.7%) and similar to the assembly of the potato genome (83.9%) (Table S2).

### Annotation of transposable elements

*De novo* annotation of repeated elements was performed with RepeatModeler version open-4-0-5 with the parameters (-engine ncbi). We identified 667 consensus repeat sequences (1.3 Mb total size) in the *N. attenuata* genome. To classify these consensus repeat sequences, additional annotation using TEclass (38) was applied for repeats that were not classified by RepeatModeler. Among all identified repeats, LTRs, DNA transposons and LINEs contributed most, representing 47.5%, 28.3% and 9.3%, respectively. The annotated repeats were used for masking repeat sequences using RepeatMasker (open-4.0.5) using parameter “-e ncbi norna”. We further re-annotated transposable elements using the *N.attenuata* repeat library for two *Nicotiana* additional genomes: *N. sylvestris* and *N. tomentosiformis* (39). To make the results comparable, we used the same approach to *de novo* identify the TE library of *S.lycopersicum* (26) and *S. tuberosum* (40) genomes.

MITEs in *Nicotiana* were annotated in two steps. First, MITE-Hunter (41) was used to find MITE families in the *N. attenuata* genome using default parameters, except “-P 0.2”. Following the manual of MITE-Hunter, the identified MITE candidate families were first subjected coverage evaluation using TARGeT. The output results were manually inspected and only the MITE families that showed even distribution of coverage were selected. Then these selected candidate MITE families were manually checked for their terminal inverted repeats (TIR) and target site duplications (TSD). In total, 15 MITE families were initially identified. We then assigned these 15 MITE families to different super-families and classes based on sequence homology to a P-MITE database. Two MITE families that showed no homology to any known MITE sequences were excluded from the downstream analysis. Second, using these 13 MITE consensus sequences as a library, we identified the copy number of each MITE family using RepeatMasker with parameters “-nolow -no_is s -s -cutoff 250”. A complete MITE sequence was defined as being no more than 3 bp shorter than the representative sequence. The multiple sequence alignment and neighbor joining tree construction of the DTT-NIC1 family were performed using ClustalW.

### Annotation of protein-coding genes

The *N. attenuata* genome was annotated using the *Nicotiana* Genome Annotation (NGA) pipeline, which employs both hint-guided (hg) Augustus (v. 2.7) (42) (hg-Augustus) and MAKER2 (v.2.28)(43), gene prediction pipelines based on genome release v1.0. For hg-Augustus gene annotation, the HMM gene model was specifically trained for *N. attenuata* using RNA-seq data from major plant tissues, and gene models were predicted using unmasked genome sequences. The repeat regions were here given less probability to be predicted as a gene, in particular when RNA-seq evidence was missing. For MAKER2 annotation pipeline, we integrated evidence of multiple protein codings from three sources: *ab initio* gene predictions, transcript evidence and protein homolog evidence. The input evidences for MAKER2 were: 1) *ab initio* gene predictors (GeneMark and Augustus) that were each trained with full-length transcripts; 2) Trinity (v. r20131110) assembled transcripts using RNA-seq data from major tissues; 3) UNIREF90 plant proteins (mapped using genewise, v. wise2-4-1); 4) six high quality plant proteomes (tomato, potato, grape, *Arabidopsis*, *Populus* and rice). The MAKER2 annotation pipeline was run on the repeat masked *N.attenuata* genome.

The predicted gene models from hg-Augustus were filtered based on their repeats contents. All genes with repeats occupying more than 50% of the gene length were removed, and genes were retained if less than 10% of their sequence matched to repeats. For genes that contained repeats occupying 10-50% of their entire gene length, we performed an additional search of the plant Refseq database using BLASTX. If the gene matched with a non-repeats homolog (e-value greater than 1e-5 and bit score greater than 200) from the plant refseq database, the predicted gene models were retained for downstream analysis. In total, 35,737 gene models from hg-Augustus predictions were retained. For MAKER2 predicted gene models, in addition to the filtering based on repeat content as described above, the gene models that had low evidence support were removed from downstream analysis (eAED <= 0.45, this cutoff was set after manual inspections on the predicted gene models). In total 33,274 gene models from MAKER2 prediction were retained. The predicted gene models from hg-Augustus and MAKER2 pipelines were then combined and overlapping gene structures were removed. After manual inspections, we found that hg-Augustus outperformed MAKER prediction when the pipelines predicted different gene structures for the same gene. Therefore, when the two pipelines predicted different gene structures for the same genome region, we retained only the hg-Augustus predicted gene models. After merging the predicted models from both hg-Augustus and MAKER2 predictions, genes with pre-mature stop codons were considered as pseudogenes and were removed from the downstream analysis. The predicted gene models were then transferred to *N. attenuata* genome release v2.0 using a custom script which first identified homologous regions using BLAST and then predicted gene structure using GeneWise. This finally resulted in 33,449 high quality gene models in the final *N. attenuata* genome, of which 74.9% were supported by at least 50 RNA-seq reads. Because gene models within a genus are usually conserved, we annotated *N. obtusifolia* and two other publicly available diploid *Nicotiana* genomes, *N. sylvestris* and *N. tomentosiformis*(39), using a homology-based annotation pipeline based on all predicted *N. attenuata* protein coding gene models. Using all predicted gene models from *N. attenuata*, we predicted protein coding gene models in *N. obtusifolia*, *N. tomentosiformis* and *N. sylvestris* using a homolog-based approach.

### RNA sequencing and data analysis

RNA was isolated from plants of the same origin and inbred generation as described above for *N. attenuata* DNA isolation. RNA was isolated using TRIZOL^®^ (Thermo Fisher Scientific) according to the instructions of the manufacturer. DNA was removed from all RNA preparations using TURBO™ DNase (Thermo Fisher Scientific) according to the manufacturer’s protocol. In total, twenty one RNA-seq libraries from different plant tissues and the same tissues under different biotic and abiotic stress treatments were first enriched for RNAs with poly-A tails and then used for RNA-seq library construction with Illumina’s TruSeq RNA sample preparation kits. The insertion sizes of the libraries are approximately 200 bp. All RNA-seq libraries were sequenced using Illumina 2000 HiSeq platform with read lengths of 50 or 101 bp and resulted in 793,785,373 paired-end Illumina reads.

The raw sequence reads were trimmed using AdapterRemoval (v1.1) with parameters “--collapse --trimns --trimqualities 2 --minlength 36”. The trimmed reads were then aligned to the *N. attenuata* genome assembly (v 2.0) using TopHat2 (v2.1.0) (44), with maximum and minimum intron size set to 50,000 and 41 base pairs (bp), respectively, estimated from the *N. attenuata* genome annotation.

To estimate the expression level of assembled genes and transcripts, all trimmed RNA-seq reads were mapped to the assembled transcripts using RSEM (v1.2.20) (45). Transcripts per million (TPM) was calculated for each transcript and gene. To exclude low-expressed genes and transcripts, only genes with TPM greater than five in at least one sample were considered as expressed. Similarly, only the transcripts with TPM greater than one in at least one sample were considered as expressed.

### Comparative genomic analysis

We assigned genes to homologous groups (HGs) using a similarity-based method. For this, we used all genes that were predicted from the 11 genomes, listed in Table 1. All-vs-all BLAST analysis was used to compare the sequence similarity of all protein coding genes, and the results were filtered based on the following criteria: e-value less than 1e-20; match length greater than 60 amino acids; sequence coverage greater than 60% and identity greater than 50%. All remaining blast results were then clustered into HGs using a Markov clustering algorithm (MLC) (46).

We constructed a phylogenetic tree for all identified HGs using an in-house developed pipeline (SI appendix).

### Evolution of nicotine biosynthesis and GCC and G-box transcription factor binding sites

Genes previously characterized as being involved in nicotine biosynthesis were retrieved from the literature and sequences were downloaded from NCBI. Phylogenetic trees for each nicotine biosynthesis gene were constructed as described above. Sequence alignment and tree structures were then manually inspected. Duplication events of nicotine biosynthetic genes were inferred from the phylogenetic tree structures as well as, when possible, from manually checking syntenic information from the tomato and potato genomes.

We extracted the GCC and G-box motif matrix from the literature (5, 27), and used FIMO (47) to detect the occurrence of these two motifs within the 2kb upstream regions of both nicotine biosynthesis genes and of their ancestral/non-root specific copies. Only the motifs with e-values less than 1e-3 were considered. Manually inspecting the positions of the annotated motif regions revealed that several motifs overlapped with annotations of TEs, such as in the upstream regions of *MPO* and *PMT*, indicating that some of these motifs may be derived from TE sequences. To test this hypothesis, we first searched GCC and G-box motif sequences within the consensus TE sequences. Overall, GCC and G-box motifs could be found in more than 54% of these TE consensus sequences. The number of GCC box and G-box motifs per kilo-base sequences in the TE consensus sequences were as high as 19 and 28, respectively. Permutation tests by randomly shuffling the positions of GCC and G-boxes 1000 times in the *N. attenuata* genome and then comparing with actual TE locations further revealed that these two motifs were significantly enriched in TE regions (*p* < 0.001). These data reveal that many TEs contain the GCC and G-box motif sequences. Next, we performed additional analyses in order to calculate the number of the GCC and G-box motifs which were derived from TE insertions that were located within the upstream regions of nicotine biosynthesis genes. For this, we extracted 150 bp sequences that included left and right flanking sequences and the motif sequence in the middle and compared these with the RepeatMasker annotated TE sequences in the *N. attenuata* genome using YASS (48), a tool designed to search for diverged sequences. To reduce false positives, only the matches that contained the expected motif sequences and had an e-value lower than 1e-5 were considered. Note that the number of GCC and G-box motifs in the nicotine biosynthesis genes that derive from TEs estimated by this approach is likely highly conservative, because this method fails to identify the corresponding homologous sequences in cases where the motif sequences and their flanking regions have diverged significantly from their ancestral TE sequences.

## Author contributions

SX and ITB conceived and coordinated the project. KG coordinated sample collections for DNA and RNA sequencing and the submission of the genome to NCBI. KG, HK, and BT coordinated the sequencing of the two genomes. HK and SX assembled the genomes. SX, TB, HT, MS and EL annotated protein coding genes in the genomes. PP and SPP annotated smRNAs in *N. attenuata*. TB and SX performed comparative genomic analysis. TB, ZL and SX analyzed RNA-seq and microarray data. SX, TB, EG and ANQ initiated and analyzed the evolution of nicotine biosynthesis and transposable elements. CK, WZ and KG validated promoter region of nicotine biosynthesis genes using Sanger sequencing, SX, EG and ITB wrote the manuscript.

The authors declare no competing financial interest.

## Acknowledgements

We acknowledge the following sources for funding: Swiss National Science Foundation (No. PEBZP3-142886 to SX), the Marie Curie Intra-European Fellowship (IEF) (No. 328935 to SX), European Research Council advanced grant ClockworkGreen (No. 293926 to ITB), DFG Exzellenzinitiative II to the University of Heidelberg (EG and ANQ) and the Max Planck Society, which provided all of the funds for the sequencing. The CoGe platform (www.genomevolution.org) is supported by NSF (IOS – 1339156 and IOS – 1444490). We thank members from the Department of Molecular Ecology for assistance with the manual curation of gene models and for scientific discussions, Dr. Sang-Gyu Kim and Dr. Matthias Erb for help with RNA sample collections, Dr. Alex Hastie from BioNano Genomics for assembling of the BioNano optical map. We thank Prof. Jonathan Gershenzon, Dr. Ewald Grosse-Wilde and Nicolas Arning for their constructive comments and suggestions on an earlier version of the manuscript.

